# Pilot study characterizing oral-to-blood microbial DNA translocation in individuals with current cocaine use disorder

**DOI:** 10.1101/2025.11.03.686400

**Authors:** Douglas Johnson, Kamala Sundararaj, Suganya Subramanian, Yaxin Qiu, Iniyan Samuel, Da Cheng, Tabinda Salman, Zhenwu Luo, Sylvia Fitting, Larry Keen, Elizabeth Call, Abhiram Maddi, William Stoops, Alicia Hartley, John E. McKinnon, Zhuang Wan, Stefano Berto, Wei Jiang

## Abstract

**Background:** Cocaine disrupts gut barriers in animal models, potentially enabling microbial translocation and inflammation in the periphery and central nervous system (CNS), but its direct role in inducing inflammation remains controversial. This study aimed to determine if the oral cavity is a source of circulating microbial DNA translocation in individuals with current cocaine use disorder (CUD).

**Results:** A cross-sectional case-control study was conducted, comparing CUD and demographically matched non-drug controls. Ten CUD (via smoking or vaping) and 24 controls provided paired saliva and blood samples. Microbial 16S rRNA V4 region was sequenced in isolated microbial DNA from saliva and plasma. Single-cell RNA sequencing (scRNAseq) was analyzed in human peripheral blood mononuclear cells. Saliva from CUD, but not plasma, exhibited reduced alpha diversity and altered beta diversity, characterized by enriched *Streptococcus* and depleted *Fusobacterium, Neisseria,* and other taxa relative to controls. Controls exhibited low to undetectable microbial translocation in plasma. By contrast, plasma displayed CUD-specific oral enrichment of several Streptococcal species and evidence of translocation into the bloodstream. *S. parasanguinis*, but not cocaine alone, induced IL-1β and TNF-α production in human primary monocytes *in vitro*. scRNAseq further revealed innate immune activation, impaired T cell function, and heightened susceptibility to infection in CUD.

**Conclusions:** This pilot study demonstrating that CUD via smoking or snorting exhibited oral microbial dysbiosis and selective oral-to-blood microbial translocation in vivo. These findings suggest that a compromised oral-to-blood barrier, rather than cocaine itself, promotes immune perturbations in CUD.

## INTRODUCTION

Cocaine, used by 5 million people in the U.S. in 2023 [1], is associated with systemic inflammation, immunodeficiency, increased susceptibility to infections, and central nervous system (CNS) abnormalities, including reduced frontal lobe grey matter and impaired cognitive function [2, 3]. The mechanisms underlying these inflammatory and immune perturbations are not fully understood.

The human oral microbiome is essential in maintaining oral health by preserving homeostasis through regulating oral immunity and metabolism [4]. Recent studies suggest that non-intravenous cocaine use causes oral microbiome dysbiosis, characterized by an enrichment of opportunistic pathogens and a reduction in beneficial commensal microorganisms [5, 6]. Oral bacterial translocation to the blood has been reported in healthy individuals after tooth brushing [7-9]. Disruption of the microbial balance may contribute to the progression of oral diseases, including periodontitis and dental caries, with implications in systemic conditions. Several bacteria translocate from the oral cavity to the circulation and have been linked to cardiovascular diseases, pre-term birth, rheumatoid arthritis, and Alzheimer’s Disease (e.g., *Streptococcus mutans, Porphyromonas gingivalis*) [10-15]. Proinflammatory microbial translocation contributes to local inflammation and facilitates the entry of bacterial products into the circulation, thereby contributing to systemic inflammation via pattern recognition receptors (PRRs), such as toll-like receptors (TLRs) [16-18].

Chronic cocaine use has been associated with heightened systemic and CNS inflammation [19-21]. Cocaine itself exhibits both immunosuppressive and immunostimulatory effects on host cells from *in vitro* [22, 23] and *in vivo* studies [24-27]. Given the ability of microbes or microbial products to enter the bloodstream following a comprised oral barrier, the potential translocation of oral microbes or their byproducts may play an indirect role in chronic cocaine use-associated persistent systemic inflammation and complications.

In this study, we report that individuals with current cocaine use disorder (CUD) via smoking or snorting exhibited altered oral microbiome by enriching cocaine-specific opportunistic pathogens (pathobionts) and depleting beneficial commensals, as well as evidence of oral-to-blood selective microbial translocation.

## METHOD

### Participants

Participants were recruited in Charleston, SC, USA. The non-drug controls were recruited via community advertisements to match the age and sex of CUD. Ten CUD and twenty-four controls who refrained from any drug use for at least one year, aged from 18 to 55 years, were enrolled. The Medical University of South Carolina (MUSC) Institutional Review Board approved this study; all participants provided written informed consent. Participants of CUD were through smoking or snorting. History or current diagnosis of psychiatric, neurological or neurodevelopmental disorders, traumatic brain injury, or recent antibiotic or probiotic usage was excluded. CUD was diagnosed through the Diagnostic and Statistical Manual of Mental Disorders, 5th edition (DSM-V), without the report of the current use of the other four drugs confirmed through chart reviews and urine 5-drug tests (Cocaine, Marijuana, Opiates, Methamphetamines, and Amphetamines, UTest Drug Testing, Framingham, Massachusetts), as described in our previous study [28]. Clinical characteristics of study participants are shown in Table 1.

**Table 1.**

| Characteristics | Controls | CUD | P value |
| --- | --- | --- | --- |
| N (M/F) | 24 (19/6) | 10 (3/7) |  |
| Age (mean y, SD) | 35.6 (10.3) | 40.3 (13.8) | NS<sup>a</sup> |
| Race (n, %) |  |  | NS<sup>c</sup> |
| White | 9 (38) | 7 (70) |  |
| Black | 5 (21) | 2 (20) |  |
| Others | 10 (41) | 1 (10) |  |

<sup>a</sup>Student's t test
<sup>c</sup>Exact test

### Microbiome analysis

The detailed microbial 16S rRNA sequencing method was shown in our previous studies [28-30]. Briefly, unstimulated saliva was collected; bacterial DNA was extracted from saliva using the QIAamp DNA Microbiome Kit (Qiagen, Germantown, MD, USA). Plasma microbial DNA was isolated from 400 μL samples using the QIAamp UCP Pathogen Mini Kit (Qiagen). Subsequently, the bacterial 16S rRNA V4 variable region was sequenced at the Microbial Analysis, Resources, and Services, University of Connecticut (Storrs, CT).

### Pro-inflammatory cytokine production in response to *S. parasanguinis*

Frozen PBMCs from four healthy donors were thawed with DNase I (20 μg/mL) [28]. *S. parasanguinis* strain F0449 (HM-808; BEI Resources, Manassas, VA) was cultured; heat-inactivated *S. parasanguinis* (1.25 × 10^6/mL, 65 °C for 30 min) was added with or without cocaine hydrochloride (5 μg/mL, Sigma-Aldrich) in the presence of GolgiStop and GolgiPlug (1 μL/mL) for 6 h. Monocyte CD14 and intracellular cytokines (Biolegend, Los Angeles, CA) were measured by flow cytometry and analyzed with FlowJo.

### Single-cell RNA sequencing (scRNAseq)

scRNAseq was performed on PBMCs from 2 CUD and 2 age-matched controls; all were white males (Singulomics Corporation, Bronx, YN). Cells were captured via 10x Genomics Chromium-X GEM-X v4, with 3′ libraries sequenced on Illumina NovaSeq X Plus (∼500 million PE150 reads/sample). Reads were processed with Cell Ranger v9.0 (GRCh38-2024-A reference, including introns) to generate gene–cell matrices; ambient RNA was removed via CellBender v0.2.0, and alignment to hg38 (gencode.v42) followed. Data integration (Seurat v5.0.0) included cell cycle analysis (CellCycleScoring), with cells retained if they had <100,000 UMIs, <20% mitochondrial reads, and no doublets (scDblFinder v1.12.0), yielding 20,017 features and 5,659 cells. Normalization, integration, and variable gene selection used SCTransform (30 PCs, resolution 0.6, UMAP); batch effects were corrected with Harmony v0.1.1. Post-RBC removal, reclustering identified 15 clusters. Cell types were annotated via FindAllMarkers, RunAzimuth, and canonical markers. Differential cell abundance was assessed with scProportionTest; DEGs (FDR < 0.05, |log2FC| > 0.3) were identified using LIBRA v1.0.0. Cell-cell communication was analyzed with CellChat v2.1.2; DEG functional annotation used scToppR v0.99.1 and clusterProfiler v4.12.2. Pathways were selected via Mann–Whitney U tests (false discovery rate [FDR] < 0.01).

### Microbiome CpG Index

To quantify the immunostimulatory potential of the gut microbiome, a CpG index was calculated for each sample based on metagenomic sequencing data. First, quality-filtered metagenomic reads were aligned to phylum-resolved reference genome databases using Bowtie2 (v2.4.5) with default sensitive alignment parameters. Uniquely mapped reads were retained to avoid cross-phylum misassignment. The CpG index reflects the net immunostimulatory capacity of microbial DNA and was computed by quantifying two classes of CpG-containing hexameric motifs: Stimulatory CpG motifs: defined as sequences matching the consensus *RRCGYY* (where R = A/G, Y = C/T), which are known to induce proinflammatory immune responses via pattern recognition receptors. Inhibitory CpG motifs: defined as sequences matching *NCCGNN* and *NNCGRN* (where N = any nucleotide), which can antagonize or suppress immune activation by stimulatory CpG motifs. For each sample, the mean frequency of stimulatory hexamers was subtracted by the mean frequency of inhibitory hexamers, and the resulting value was normalized per kilobase of effectively mapped sequence to account for differences in sequencing depth and genome length. This calculation followed the computational framework established by Lundberg et al. [31], ensuring comparability with prior immunomicrobiome studies. To generate phylum-level CpG indices, taxon-specific CpG values were first estimated for each identified microbial feature within a given phylum. These values were then aggregated by computing a weighted average across constituent taxa, with weights proportional to their relative sequence abundance in the sample. This abundance-weighted aggregation yields a representative CpG index for each bacterial phylum, reflecting the overall immunostimulatory profile of that lineage within the gut microbial community.

### Statistical Analysis

OTU counts were normalized to relative abundances, and taxa were aggregated at phylum, class, order, family, genus, and species levels. To evaluate for a significant difference in composition (i.e., whether the groups occupy distinct regions in multivariate space, as hinted by the plot), a permutational multivariate analysis of variance (PERMANOVA) was conducted on the full Bray-Curtis distance matrix. Group differences in abundances were assessed using the non-parametric Mann–Whitney U test in QIIME 1, with FDR correction. α-diversity within samples was calculated using the phyloseq package in R and compared by the Wilcoxon rank sum test. Differences in overall microbiome composition between groups were evaluated by permutational multivariate analysis of variance (Adonis function, vegan package in R), with P values adjusted by the Benjamini–Hochberg FDR method. Statistical significance was defined as P < 0.05. Analyses were performed using GraphPad Prism (version 10.4.1, GraphPad Software, Boston, MA, USA).

## RESULTS

### CUD-associated microbiome dysbiosis in saliva but not in plasma

In saliva, α-diversity (Simpson Index) was significantly reduced at the phylum (p < 0.0001), genus (p < 0.0001), and species levels (p < 0.05) (Figure 1A–1C), indicating decreased oral bacterial richness in CUD. In plasma, no difference of α-diversity was determined between the two study groups (P > 0.05). β-diversity (Bray–Curtis PCoA) revealed significant group differences in saliva at all levels, as well as plasma at the genus level (p = 0.02) but not at the phylum or species level (p > 0.05) (Figure 1D–1F). These findings demonstrate that CUD significantly disrupts the oral microbial community, with minimal changes in circulating microbial diversity.

**Figure 1.**
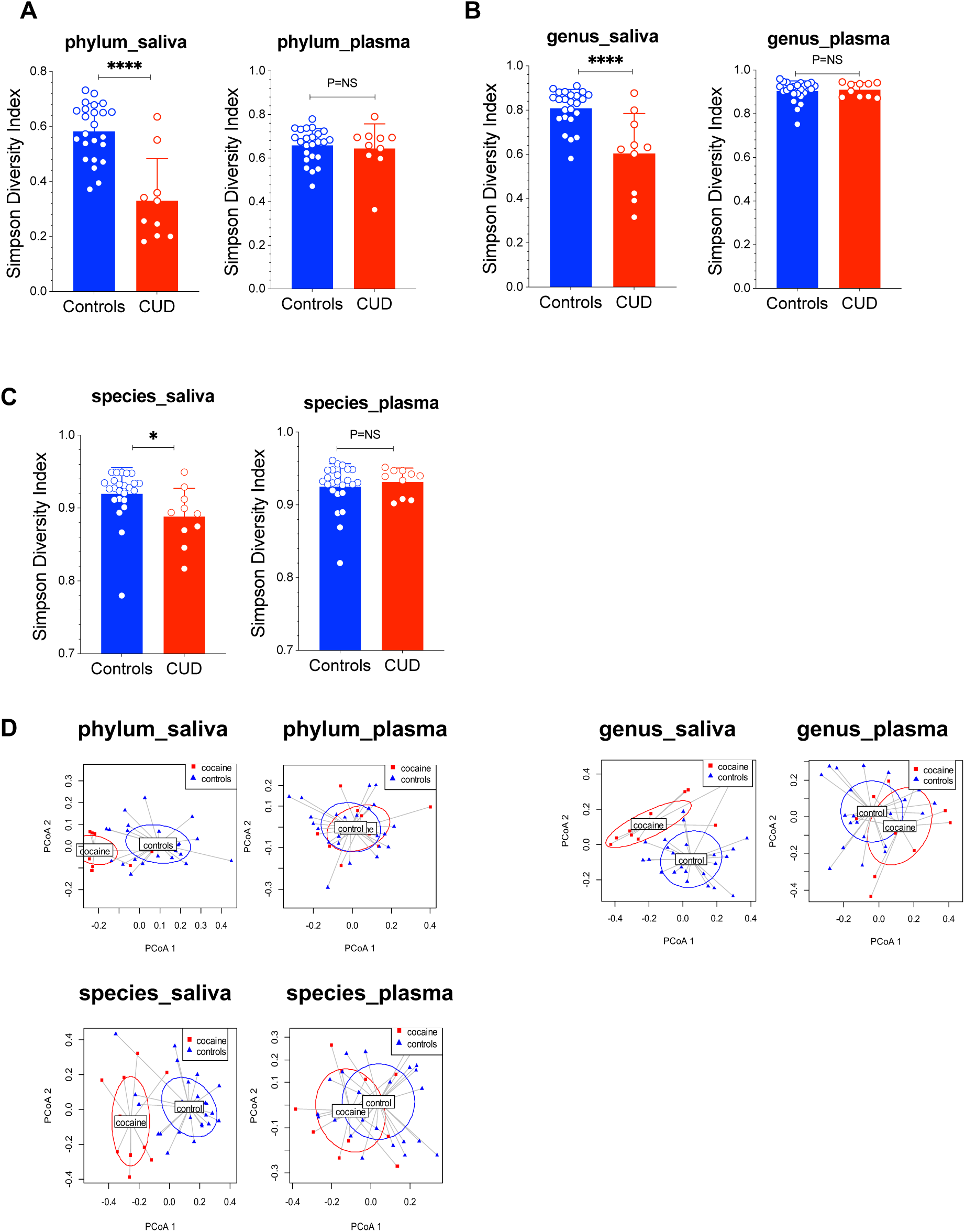
Oral but not blood microbial dysbiosis in CUD. (A-C) Saliva and plasma microbiome α-diversity (mean ± SD, non-parametric MannWhitney tests) and (D-F) β-diversity in CUD and controls. **** p<0.0001, NS: P > 0.05. Each point represents an individual sample, with horizontal lines indicating group means. A permutational multivariate analysis of variance (PERMANOVA) was applied to β-diversity on the full Bray-Curtis distance matrix.

### Relative abundance of bacteria in saliva and plasma

We next assessed the enrichment of taxa in CUD versus controls. Overall, more taxa were detected with low abundance in the plasma compared to saliva at each levels, suggesting plasma microbial translocation from various sources besides the oral cavity (Figure 2A-2B, supplemental Figures 1-2). At the phylum level (fold changes in CUD versus controls, supplemental Figure 1A), CUD saliva was enriched for *Firmicutes* and *Actinobacteria* and depleted of *Fusobacteria*, *Bacteroidetes,* and *Proteobacteria*, whereas CUD plasma showed higher *Proteobacteria, Actinobacteria, Firmicutes,* and lower *Nitrospirae,* and others. Notably, heatmap analysis revealed *Streptococcus* as the predominant oral genus translocated into plasma in CUD but not controls, despite being orally present in both groups (Figure 2A; supplemental Figures 1B and 2A). At the genus level, CUD saliva was enriched for *Streptococcus* and depleted of *Fusobacteria*, *Granulicatella, Heamophilus, Leptotrichia, Neisseria, Porphyromonas,* and *Proteobacteria*, whereas CUD plasma showed higher *Streptococcus* and *Neisseria* and lower *Leptotrichia* and *Porphyromonas* (Figure 2A). At the species level (Figure 2B, supplemental Figure 2B), profiling showed higher *Streptococcus species* (e.g., *S. australis*, *S. parasanguinis*, *S. vestibutaris,* and *S. thermophilus*) and *Veillonella dispar*, with lower species including *Neisseria mucosa, N. cinerea, N. flavescens, Prevotella melaninogenica, Granulicatella adiacens, Actinobacillus porcinus*, and *Haemophilus parainfluenzae* in saliva of CUD. Among the four Streptococcal species increased in saliva of CUD, *S. australis* and *S. parasanguinis* increased in plasma of CUD versus controls.

**Figure 2.**
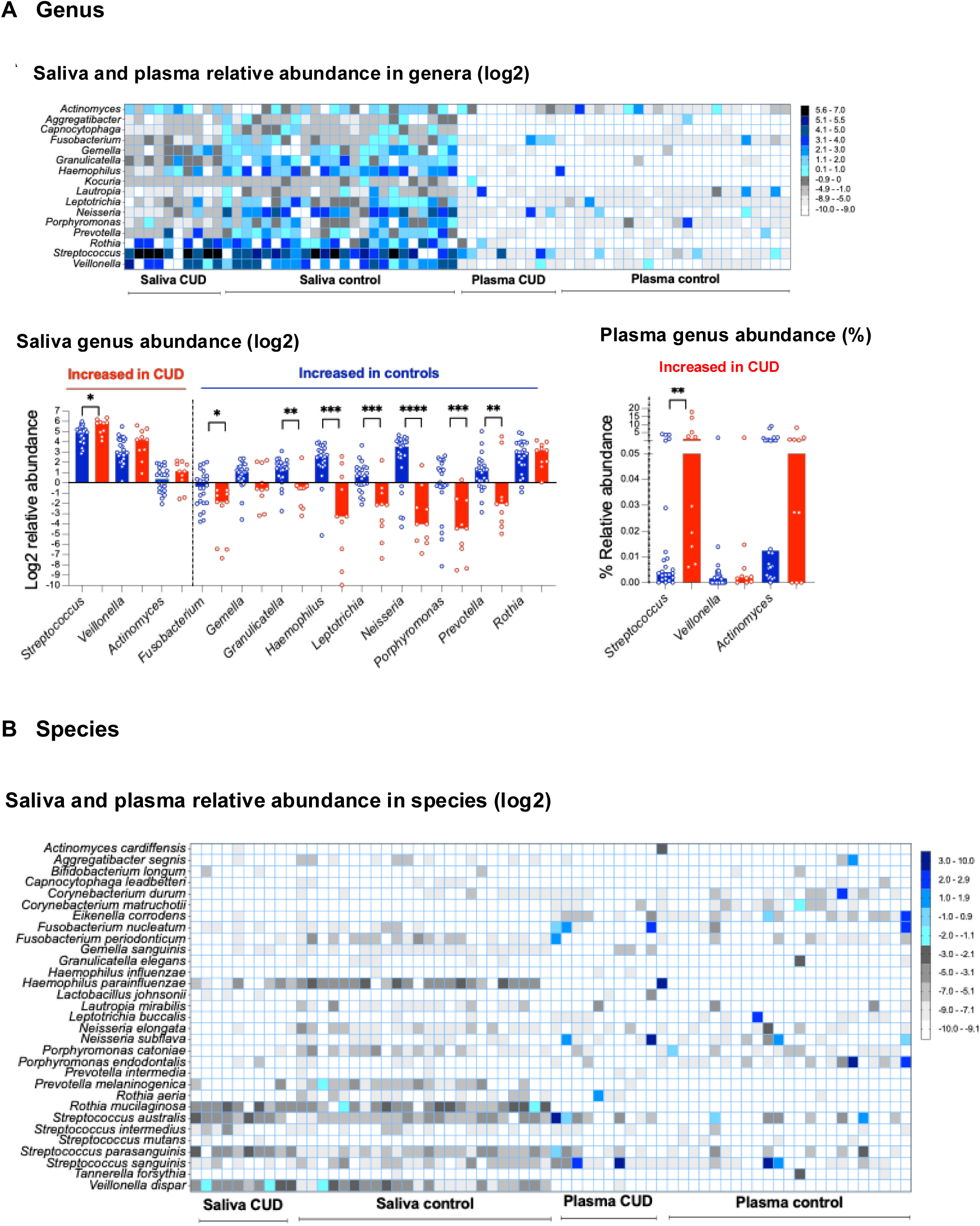
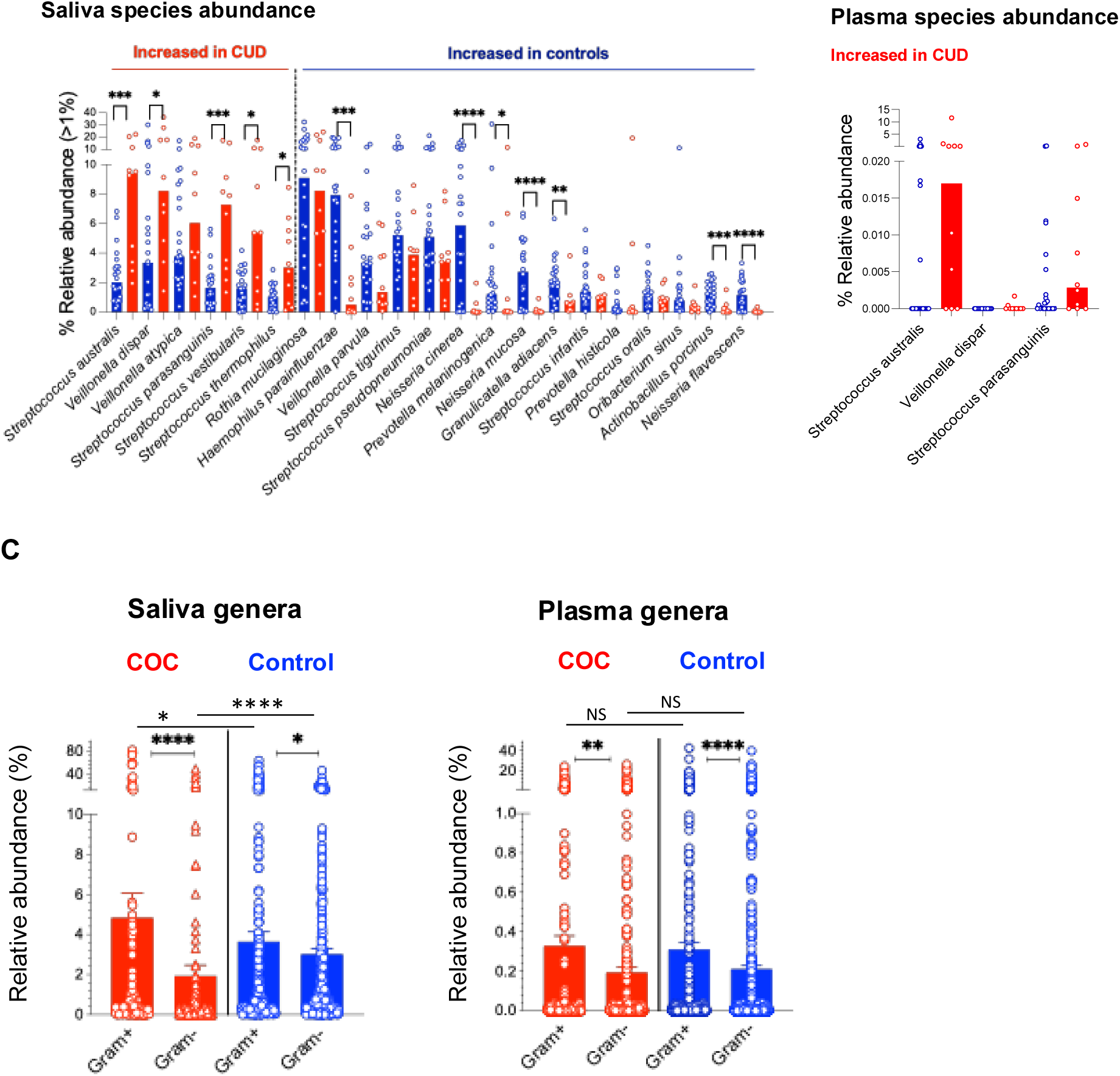
Comparative analysis of microbial abundance in saliva and plasma samples from CUD versus controls. (A-B) Heatmap (median relative abundance, log2 of percentages) and dot plots (median relative abundance, log2 of percentages) analysis of shared microbial genera and species in saliva and plasma samples. Shared taxa in saliva were selected from the top-enriched genera or species in any group. (C) Relative abundance of gram-positive and gram-negative genera in saliva and plasma. Mean ± SEM.

### Increased abundance of saliva gram-positive bacteria in CUD versus controls

We next evalute the enrichment of gram-positive and negative bacteria (Figure 2C), and found that overall more gram-positive versus gram-negative in both saliva and plasma from both groups; saliva but not plasma from CUD exhibited further increased abundance of gram-positive bacteria compared to controls. These findings suggest a relatively consistent oral-to-blood microbial shift in CUD, whereas in controls, the blood microbiome appears to derive more from other sources (e.g., gut) besides the oral cavity.

### Oral-to-blood microbiome DNA translocation

To futher evaluate oral-to-blood microbial translocation, we analyzed shared genera and species in both sampling sites and compared between the two study groups (Figures 3A-3B, median ± 95% CI). Overall plasma microbial translocation was significantly lower in controls than in CUD patients, reflecting an intact gut barrier in healthy individuals but not in those with CUD. Notably, CUD exhibited mainly increased *Streptococcus* in both sites, especially in the plasma, compared to controls. At the species level (Figure 3B), *S. sanguinis*, *S. australis*, and *S. parasanguinis* were enriched in both saliva and plasma of CUD, consistent with oral-to-blood translocation. In contrast, controls exhibited higher abundances of *Neisseria* and *Haemophilus*, hallmarks of a balanced oral microbiome [32], while CUD showed a shift toward opportunistic pathogenic taxa.

**Figure 3.**
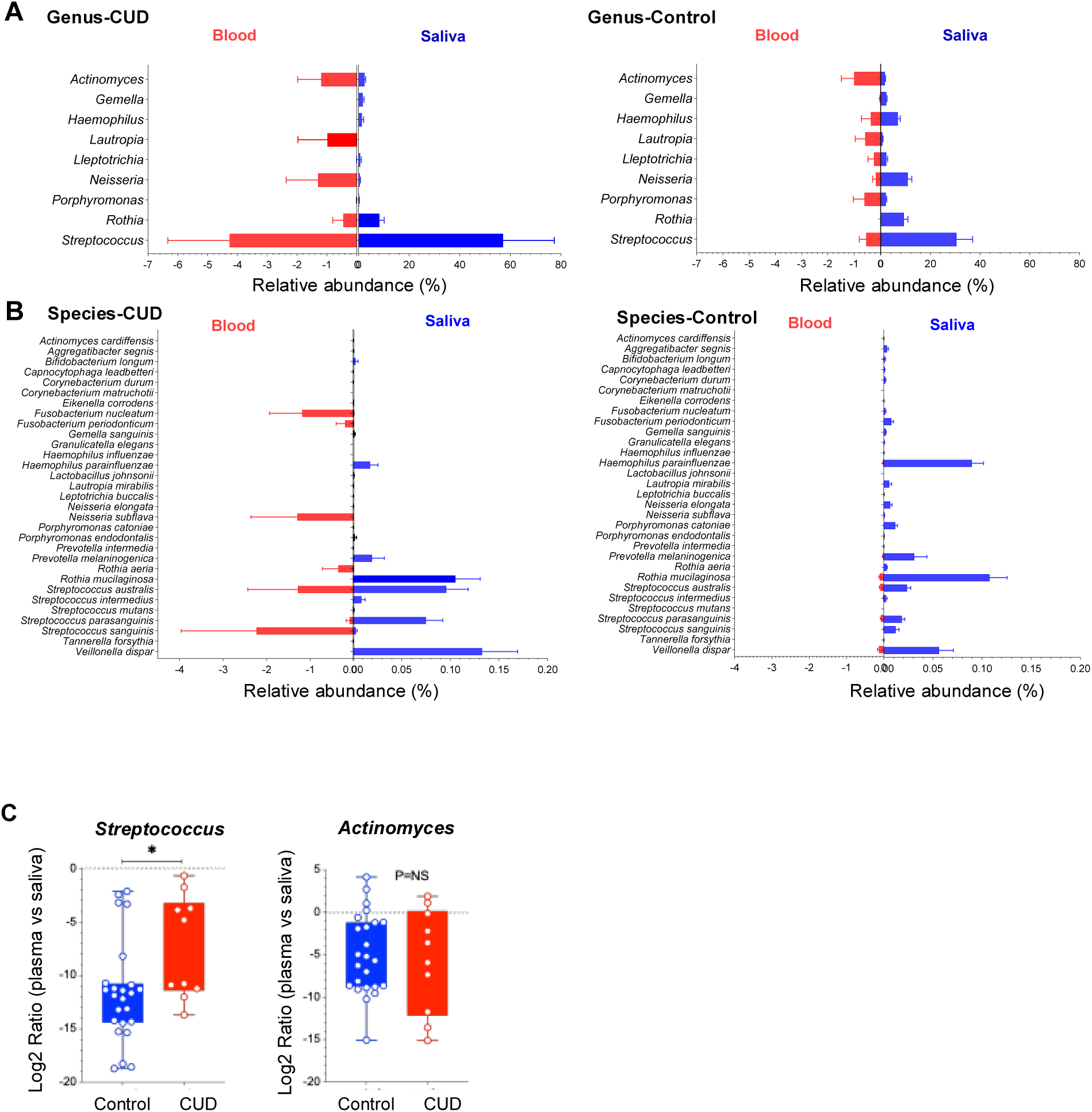
Oral-to-blood microbiome consistency. (A-B) Relative abundance of bacterial genera (A) and species (B) in saliva and plasma samples of CUD and controls, emphasizing potential oral-to-blood tranlocation. Data is shown as median ± 95% CI; taxa was selected for detecting in both sites. (C) Plasma-to-saliva ratios (log2) of *Streptococcus* and *Actinomyces* between the two study groups.

Focusing on oral-associated taxa [33], *Streptococcus* and *Actinomyces* were increased in both saliva and plasma of CUD (Figure 2A and Figure 3A). The plasma-to-saliva ratio of *Streptococcus*, but not *Actinomyces*, was elevated in CUD compared with controls (Figure 3C), suggesting differential translocation potential. In controls, *Streptococcus*, *Neisseria*, *Haemophilus*, and *Rothia* remained largely restricted to saliva and rarely detected in plasma. These findings indicate selective oral-to-blood microbial DNA translocation in CUD, reflecting compromised barrier function and dysbiosis.

Host cell interactions with various microbes lead to the production of pro-inflammatory cytokines [16]. To verify whether *S. parasanguinis* exhibits proinflammatory activity, we cultured PBMCs with heat-inactivated *S. parasanguinis* and found a noticeable increase in the production of TNF-α and IL-1β (Figure 4). However, cocaine did not impact the cytokine production (Figure 4). These findings suggest that cocaine may promote systemic inflammation indirectly by enriching and translocating proinflammatory microbiome elements, rather than acting as a direct inducer.

**Figure 4.**
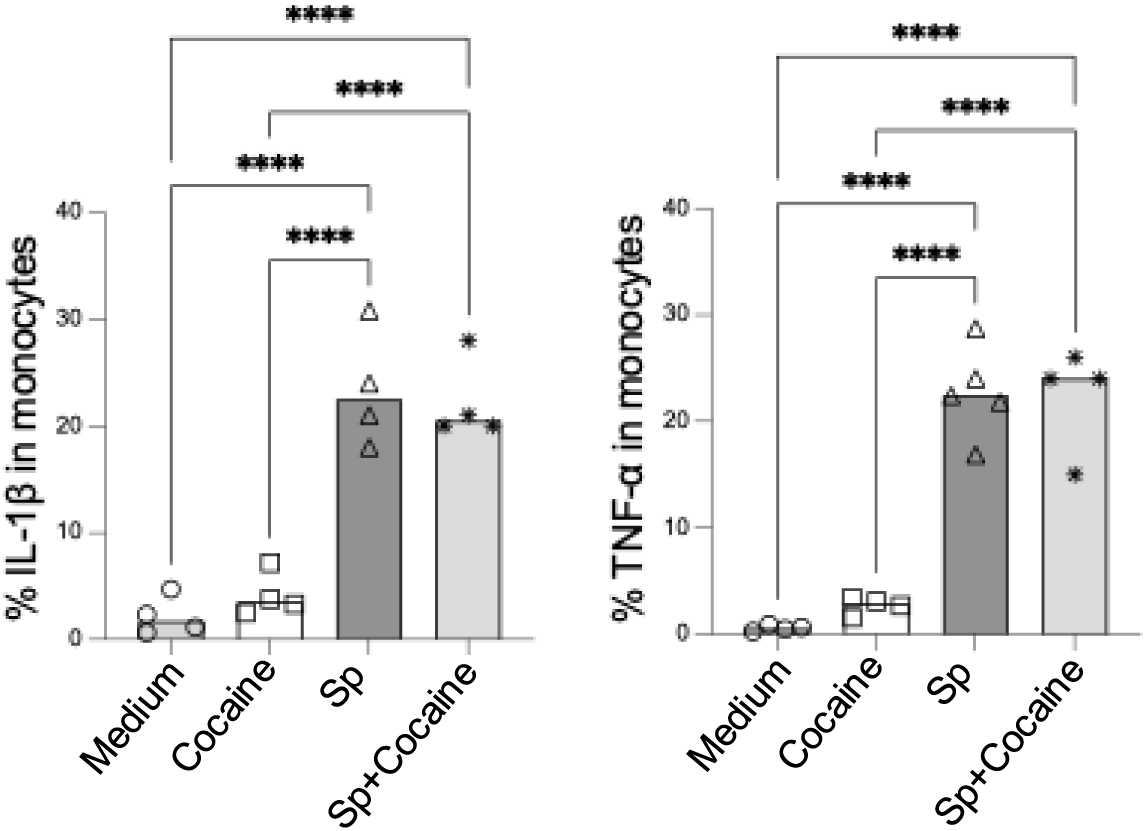
*S. parasanguinis* promotes proinflammatory cytokine production *in vitro*. PBMCs isolated from four healthy donors were stimulated with *S. parasanguinis* (SP), cocaine, or a combination of SP and cocaine. The production of pro-inflammatory cytokines TNF-α and IL-1β was measured using ELISA. Data are presented as mean ± SD, and statistical significance is indicated as follows: **p < 0.01; ****p < 0.0001.

### Reduced CpG index in oral microbiome phyla of CUD

The CpG index, measuring bacterial DNA’s net immunostimulatory potential, was calculated for the five dominant phyla (Firmicutes, Bacteroidetes, Proteobacteria, Actinobacteria, Fusobacteria) in each sample (Figures 5A-5B). All phyla had positive CpG indices (indicating oral microbiome immunostimulatory potential), with plasma microbiome showing unexpectedly higher CpG indices than saliva, suggesting that CUD is associated with lower immunostimulatory potential of the oral microbiome. Aggregate phylum-level CpG indices were significantly lower in CUD than controls. Oral phyla of controls (Firmicutes, Bacteroidota, Actinobacteriota, Proteobacteria) had CpG indices clustering near the neutral range indicating relative genomic stability, while CUD had significantly reduced CpG indices indicating oral microbial CpG suppression. The CpG index reduction was most pronounced in *Firmicutes* and *Actinobacteriota*, phyla enriched in opportunistic taxa (e.g., *Streptococcus, Actinomyces*) with oral-to-blood translocation evidence. This suggests CUD-associated oral dysbiosis favors microbes with greater CpG depletion linked to host–pathogen interactions and immune evasion, indicating CUD alters oral microbiome taxonomic composition and genomic CpG content.

**Figure 5.**
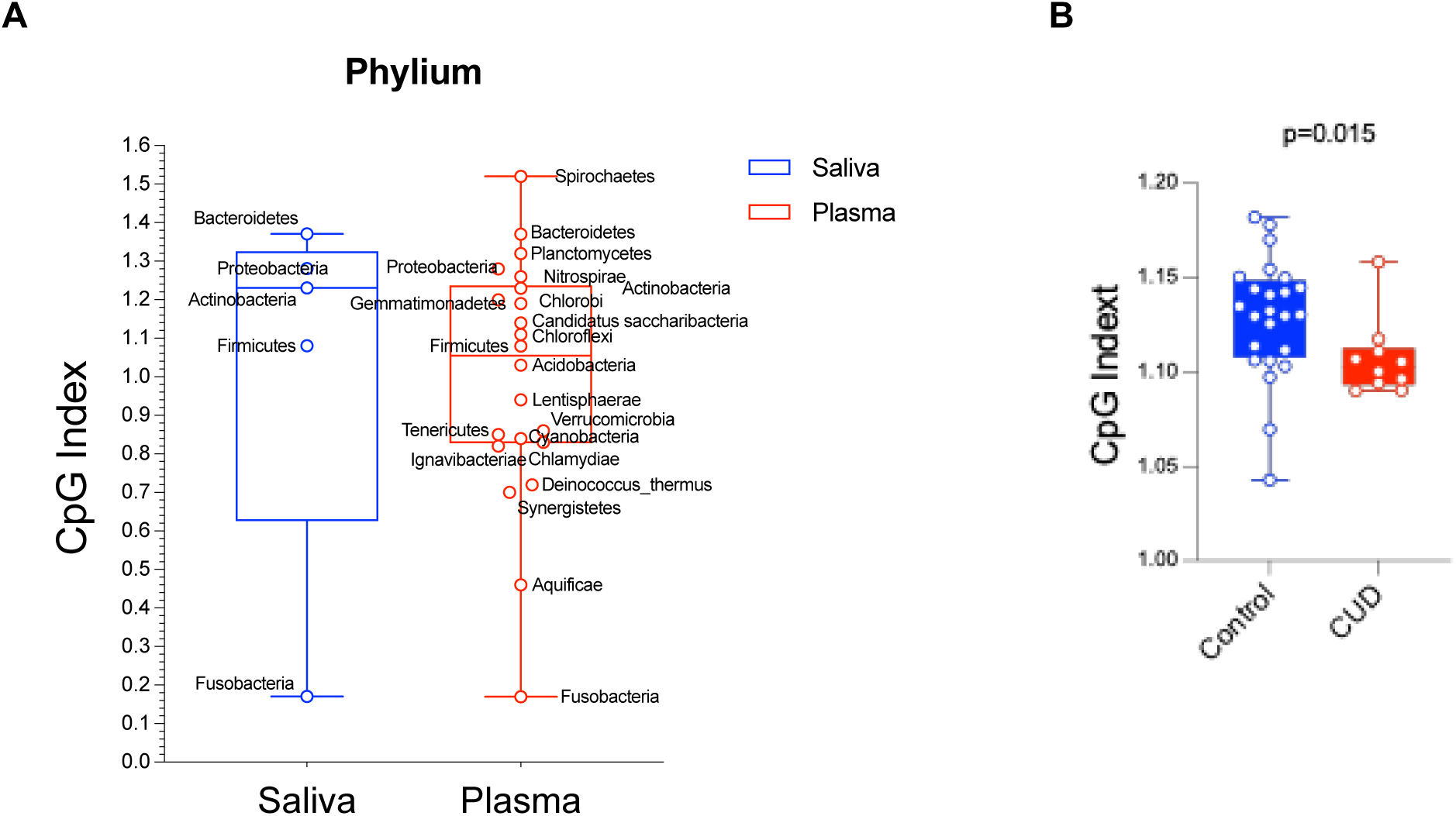
Distribution of CpG indices at the phylum level for CUD (red) and control (blue) groups. (A) CpG indices in each genera from saliva and plasma. (B) CpG indix comparisons in the two study groups. Comparisons of CpG index distributions in CUD were lower than controls. Data are presented as boxplots showing median, interquartile range (IQR), and individual sample distributions. Phylum-specific analyses revealed the most pronounced reductions in Firmicutes (p = 0.002) and Bacteroidetes (p = 0.004) in CUD, which may contribute to altered immune responses in the oral cavity. No significant correlations were found between CpG index and cocaine use duration or frequency (Spearman’s ρ < 0.2, p > 0.1).

scRNAseq profiling reveals distinct circulating immune perturbations in CUD versus controls scRNAseq of PBMCs identified 12 immune populations (e.g., CD4+ and CD8+ T cell subsets, CD14+/CD16+ monocytes, DCs, B cells, NK cells, platelets) in CUD and controls (Figure 6A-6E). CUD showed immune composition shifts: increased NK cells, total B cells, and platelets; reduced CD4+ and CD8+ T cell subsets and naive B cells (Figure 6C-6D; supplemental Figure 3). Differential gene expression (DGE) was most prominent in CD8+ effector T cells (1,504 DEGs),

**Figure 6.**
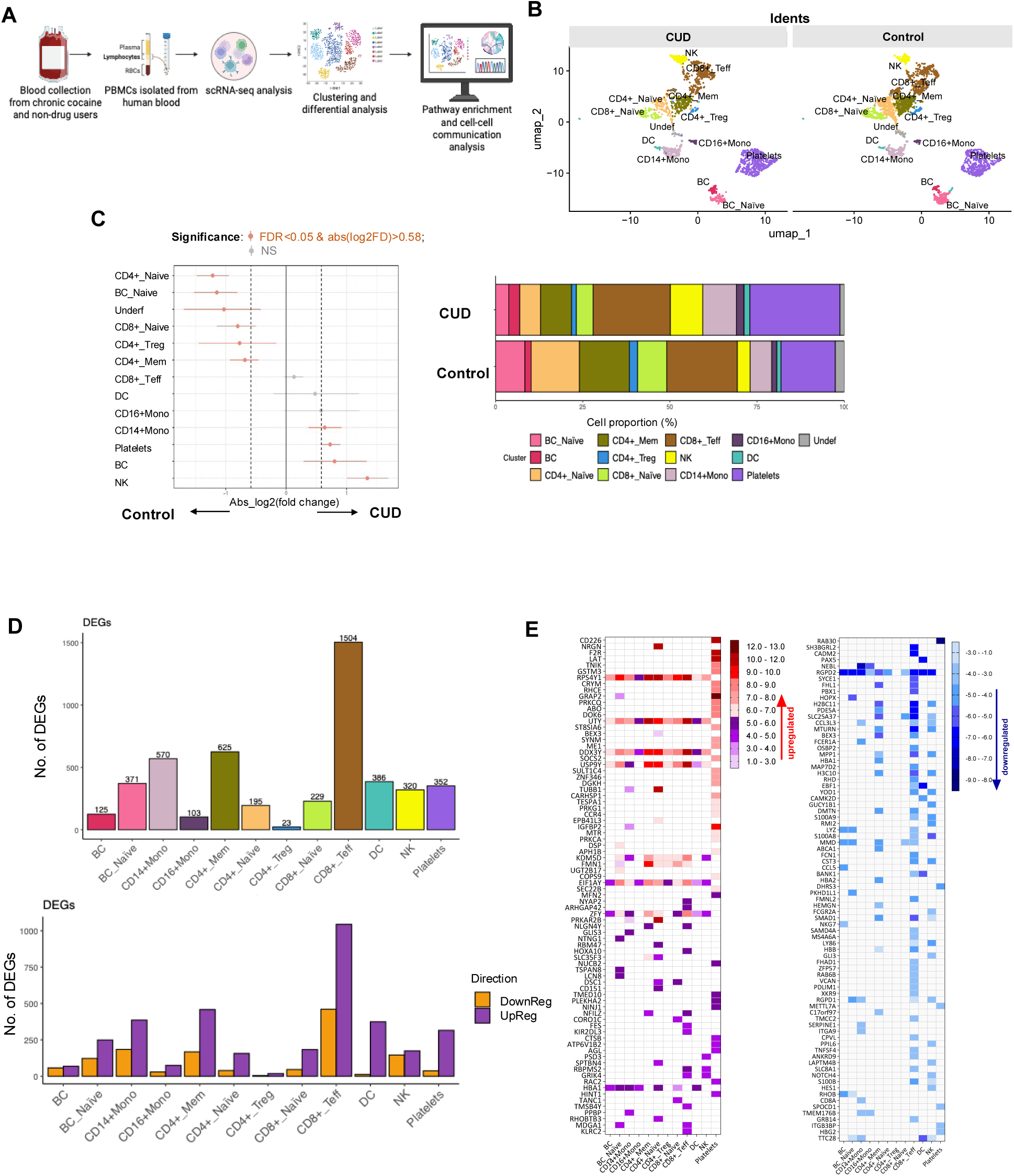
scRNAseq of PBMCs from CUD and controls. (A) Workflow schematic of scRNA-seq analysis. (B) UMAP plots showing distinct clustering of immune cell populations in CUD versus controls. (C) Differential cell-type composition, with significant shifts in NK, B cells, monocytes, and platelets, alongside stacked bar graphs of relative proportions. (D) Bar plots of DEGs across immune cell populations, showing both total numbers and the numbers of up- and down-regulated DEGs. CD8+ effector T cells, monocytes, and memory CD4+ T cells showed the most significant transcriptional changes. (E) Heatmap of representative DEGs by cell type, with red indicating upregulation and blue indicating downregulation, based on the top means of 80 up-and 80 down-regulated genes in CUD versus controls in any cell type.

CD4+ memory T cells (625), CD14+ monocytes (570), and DCs (386) (Figure 6D). Most DEGs were upregulated, including genes related to immune activation and function, such as RPS4Y1, UTY, and DDX3Y, in lymphocytes and monocytes but not platelets, NRGN (CD4+ naive T cells, adaptive neuroimmune response), and HBA1 (multiple cells, oxidative stress compensation) (Figure 6E and 7A). Downregulated DEGs included RGPD2 (broad cell types, cellular processes), NEBL (monocytes, cytoskeletal function), and CCL3L3, FCER1A, and CD8A (monocytes, innate immune suppression) (Figure 6E and 7A).

**Figure 7.**
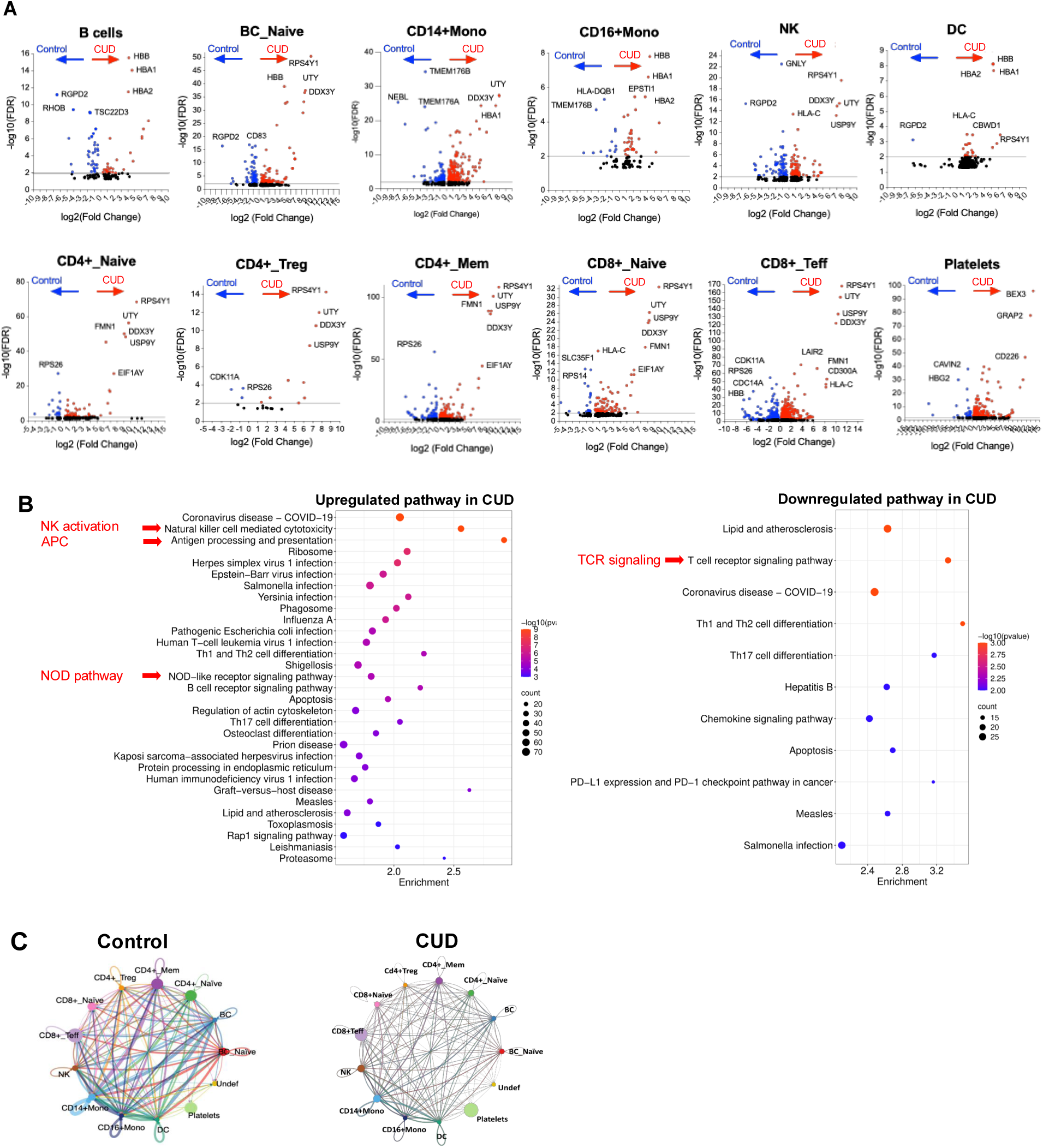
Differential gene expression, pathways, and cell-cell communication in CUD versus controls. (A) Volcano plots of representative immune subsets show significantly upregulated (red, ratio of CUD versus controls) and downregulated (blue, ratio of CUD versus controls) genes in CUD versus controls, with top markers labeled. Above the grey line indicates an adjusted p-value < 0.01. (B) Pathway enrichment analysis highlights upregulation of NK activation, antigen presentation, NOD-like signaling, and infection-related pathways, alongside downregulation of TCR signaling, Th differentiation, chemokine signaling, and PD-1 checkpoint pathways. (C) Chord diagrams illustrate predicted ligand–receptor interactions, with controls showing balanced networks and CUD exhibiting altered connectivity and disrupted immune communication.

Pathway analysis revealed CUD upregulated innate immunity (NOD-like receptor signaling, NK activation, antigen processing) and pathogen response pathways, but downregulated adaptive immunity (TCR signaling, Th cell differentiation, chemokine signaling) (Figure 7B). Cell-cell communication in CUD showed reduced network density and connectivity (especially T cells and monocytes) compared to controls (Figure 7C), indicating impaired intercellular coordination. These findings support CUD-driven innate immune activation, adaptive immune suppression, and immune dysregulation, likely linked to microbial translocation (e.g., S. parasanguinis) rather than cocaine directly.

## DISCUSSION

In this study, CUD was characterized by oral microbial dysbiosis, reflecting the expansion of opportunistic oral pathogens (e.g., *Streptococcus spp*.). While several species were consistently enriched in both saliva and plasma when comparing CUD versus controls, the data suggest that plasma microbial translocation in CUD originates at least partially from the oral cavity. Controls exhibited significantly low to undetectable levels of microbial translocation. However, CUD-specific oral-to-blood translocation may be diluted within the broader physiological background of microbial translocation from other compartments, such as the gut, which was evident in controls.

Chronic cocaine use is strongly associated with oral health deterioration, including periodontitis [34]. Reduced salivary flow, a common consequence of cocaine use, diminishes pathogen clearance and contributes to oral dysbiosis [35]. Cocaine also exerts immunomodulatory effects that favor colonization by opportunistic pathogens [34], increasing susceptibility to infections [36, 37]. Studies conflict on the direct impact of cocaine on anti- or pro-inflammatory responses, ranging from enhanced proinflammatory responses to inhibition of myeloid cell activation [22-27]. In humans, acute cocaine administration suppresses peripheral inflammation, reducing monocytic TNF-α and IL-6 production [38]. By contrast, chronic cocaine use is linked to dysregulated systemic inflammation [39] and neuroinflammation, characterized by microglial activation and neuropathology in both animal models and humans [40, 41]. These effects may arise from direct neurotoxicity on glial cells [40] as well as indirect mechanisms. Prolonged use disrupts mucosal barrier integrity and alters microbiota composition, promoting microbial translocation into circulation. While some studies suggest cocaine may impair epithelial barriers directly [19, 42], others indicate that barrier disruption is not solely cocaine-mediated [22]. Instead, microbiota-driven dysbiosis is likely to contributes to barrier dysfunction. Thus, we propose that microbial shifts, rather than cocaine itself, drive host immune activation, and persistent inflammation.

Cocaine use induces a compromised barrier and alters gut microbiomes in previous human and animal studies [19, 43, 44]. Although most microbiome research has focused on the gut, our data show that in CUD, the plasma microbiome partially mirrors that of saliva. These results implicate the oral cavity exposed during smoking or snorting as a source of microbial translocation into the bloodstream in CUD.

The CpG index, a measure of the immunostimulatory potential of bacterial DNA, was calculated at the phylum level using representative reference genomes for dominant taxa within each phylum, weighted by their relative abundance in the metagenome. Lower CpG reflects lower immunostimulatory potential. Controls displayed a wider range of CpG values, characterized by a higher median index and greater variability, consistent with a more diverse and potentially immunostimulatory microbial milieu. In contrast, individuals with CUD displayed clustered, lower CpG index values and fewer high-index outliers, indicating reduced overall immunostimulatory potential consistent with a direct inhibitory effect of cocaine within the oral cavity. Nevertheless, the elevated abundance of oral bacterial taxa such as *Streptococcus spp*. in CUD is inherently proinflammatory as a general microbial trait; consequently, these organisms may drive chronic systemic inflammation through microbial translocation into the circulation. Collectively, these findings support the hypothesis that cocaine use-associated impairment of oral barriers, rather than a cocaine-induced shift in the oral microbiome toward a proinflammatory community, constitutes a central pathogenic mechanism enabling oral microbiome translocation.

To our knowledge, this is among the first human study to provide evidence of such a compromised barrier in CUD, marking a significant step in understanding the systemic consequences of CUD via oral health disruptions. However, we cannot entirely exclude the possibility that oral microbiome DNA translocates to the blood via the gut. Nonetheless, multiple pieces of evidence from other colleagues support the translocation of oral-to-brain rather than oral-to-gut in CUD: 1) human studies using paired samples from different sites, revealed that the predominant colonization of *Streptococcus* species; for example, *S. parasanguinis* exhibits 100-1000-fold higher copies per ng of microbial DNA in the saliva versus stool [33, 45]; and 2) clinical evidence of bacteria translocation from the oral cavity to the system contributing to the development of preterm birth, stroke, and metabolic disorders [12, 46, 47]. Our results highlight that chronic cocaine exposure via smoking or snorting destabilizes the oral ecosystem, enabling persistent oral-to-blood translocation, which may contribute to CUD-associated complications, including elevated innate immunity and systemic inflammation, immune perturbations, and susceptibility to infections.

Oral *Streptococci* have a dual nature. While typically harmless (or even protective) in the oral cavity, if they or their byproducts gain access to systemic body sites, they can cause disease. For example, many oral *Streptococci* express surface proteins that bind extracellular matrix components (collagen, fibrin) and platelet receptors, facilitating their attachment at sites like damaged endocardium or blood vessels and contributing to systemic diseases [48-51]. However, these cases occur more frequently in immunocompromised individuals. Certain oral *Streptococcus* strains possess immune-evasive and proinflammatory properties that facilitate both colonization and pathogenicity. Traits such as adherence, biofilm formation, and immune modulation, while critical for oral biofilm stability, also enable invasive potential when bacteria are displaced [52, 53]. Chronic oral dysbiosis, as in periodontitis, exacerbates this risk by increasing bacterial burden and weakening mucosal integrity. Periodontitis-associated *Streptococcus* species can persist within inflamed gingival tissues, provoking sustained systemic immune activation or intermittently translocating into circulation [54, 55]. Thus, *Streptococcus* species play a context-dependent role, as essential contributors to oral homeostasis under balanced conditions, but potential drivers of systemic inflammation and infection when homeostasis is disrupted. Their capacity to modulate immune responses locally and systemically underscores their importance in linking oral health to overall health. Moreover, the potential for oral taxa (including *Streptococcus* spp.) to influence tryptophan metabolism and CNS signaling via microbial neurotransmitter pathways, and targeted metabolomics should be explored in future work.

In saliva, opportunistic taxa such as *Streptococcus* were enriched, whereas commensals *Neisseria, Prevotella,* and *Haemophilus* were depleted, consistent with oral dysbiosis. Among these changes in saliva, several were also observed in plasma. The detection of specific *Streptococcus* species in both saliva and plasma suggests oral-to-blood microbial DNA translocation in CUD (Figure 3B). Notably, *Neisseria* was a highly enriched genus in control saliva (controls versus CUD, Figures 2A and 3A), whereas *N. subflava* was detected exclusively in the blood of CUD but low in controls (Figure 3B); *F. periodonticum, F. nucleatum* and *R. aeria* were identified only in the blood of CUD (Figure 3B), consistent with a compromised oral–blood barrier. Despite site-specific differences, overall microbial enrichment patterns were similar between groups, suggesting selective and microbiome-specific translocation that warrants further investigation.

It is important to note that our observed microbial DNA translocation (consistent with low-biomass plasma levels) and that bacterial products (DNA fragments, cell wall components) likely drive inflammation rather than whole live bacteria (no bacteremia) [56]; no clinical infection was observed in all study participants during the study visit. Instead, we propose that bacterial byproducts, such as DNA fragments, bacterial cell wall products (i.e., LPS and PGN), and other microbial antigens (i.e., microbial metabolites), show different abilities to induce proinflammatory cytokine production [30] via entering the bloodstream persistently and contributing to chronic systemic inflammation [57, 58], deserving further investigations. This perspective aligns with CUD-mediated systemic inflammation and pathology indirectly via oral-to-blood microbial translocation or entering through the lymphatic system [59].

Building on prior reports of immune dysregulation and immunodeficiency, impaired adaptive immunity, and elevated rates of cardiovascular, infectious, and neuropsychiatric comorbidities in CUD [39-41, 60], scRNA-seq revealed heightened activation of innate immune pathways (e.g., NOD-like receptor signaling, NK cell cytotoxicity) and increased infection susceptibility, accompanied by broad suppression of T-cell receptor signaling in CUD, indicating innate hyperactivation coupled with impaired adaptive immunity. Predicted disruption of cell–cell communication networks further indicated compromised immune coordination.

While our study is the first to illuminate the potential for oral-to-blood microbial translocation in CUD, it is not without limitations. 16S rRNA V4 sequencing, employed herein, lacks precise species-level resolution, metagenomic shotgun sequencing or full-length 16S RNA sequencing represents a more advanced approach for future investigations. Notably, *in vitro* experiments demonstrating cocaine-induced growth of *S. parasanguinis* align with our *in vivo* human microbiome findings at the species level, mitigating this limitation in part. Missing data on oral hygiene, cigarette smoking frequency, and diet is a limitation. Additionally, scRNA-seq results indicate transcriptional and pathway-level changes, not direct functional assays, the small sample size of scRNA-seq data restricts the scope of conclusions we can draw, and ex vivo functional validation was partially addressed through monocyte cytokine assays, but remains limited. Dental exams were not included in the IRB-approved protocol due to logistical constraints (recruitment in a non-dental setting); future studies should potentially integrat periodontal indices or imaging. Finally, contamination risk in plasma microbiome studies needs to be aware and controlled (i.e., adding water controls). To fully elucidate the temporal dynamics of microbial translocation and its downstream health impacts in CUD, causal and longitudinal studies are warranted.

## CONCLUSIONS

This study provides initial human in vivo evidence of selective oral-to-blood microbial DNA translocation in CUD, rather than only descriptive dysbiosis. The findings are hypothesis-generating and exploratory. The selective enrichment of *S. parasanguinis* growth suggests cocaine promotes the expansion of pathobionts with potential systemic sequelae. Single-cell transcriptomic analyses further revealed a signature of innate immune activation, heightened infection susceptibility, and compromised T-cell-mediated adaptive immune responses, supporting a model wherein translocated microbial products sustain immune dysregulation. Notably, to our knowledgement, this is among the first *in vivo* study to link CUD-associated oral barrier dysfunction to systemic immune perturbations. These findings highlight the oral cavity as a key yet underappreciated driver of systemic immune dysfunctions in CUD, bridging local microbial alterations to broader systemic disturbances. Future research should investigate whether targeted microbiome interventions can restore oral barrier function, rebalance host–microbe crosstalk, and ultimately alleviate CUD-associated systemic pathogenesis.

## AUTHOR CONTRIBUTIONS

D.J., K.S. I.S., A.H., and Z.L., performed experiments. D.J. and Y.Q. wrote the manuscript. Z.W., W. X., D.J., S.S., and W.J. analyzed data. E.C., D. C., L.K., J.E.M., W.S., T.S., S.B., S.F., A.M., and W.J. revised the manuscript.

## ACKOWLEDGEMENTS AND FUNDINGS

This work was supported by grants from the National Institutes on Drug Abuse R01DA059854 (Jiang), R01DA059538 (Jiang), R01DA055523 (Fitting & Jiang), 2R15DA052886-02 (Keen), 2026 Brain & Behavior Research Foundation (BBRF) Distinguished Investigator Award (34152, WJ), and by the Odyssey Pre-doctoral Fellowship from the Medical University of South Carolina College of Graduate Studies, the CNDD Genomics and Bioinformatics Core at MUSC (NIH grant P20GM148302), and by the Biorepository & Tissue Analysis Shared Resource, Hollings Cancer Center, Medical University of South Carolina (P30 CA138313).

## HUMAN ETHICS AND CONSENT

The Medical University of South Carolina (MUSC) Institutional Review Board approved this study; all participants provided written informed consent.

Clinical trial number: not applicable.

## DATA AVAILABILITY STATEMENT

The NCBI Gene Expression Omnibus (GSE304577) accession number for the single-cell RNA-seq data is available. Single-cell data is available through a shiny app: https://bioinformatics-musc.shinyapps.io/JiangLab_COCAINE_study/.

Custom R codes and data to support the analysis, visualizations, and functional enrichments are available at https://github.com/BioinformaticsMUSC/JiangLab_CUDAINEproject.

## Figure legends

**Supplemental Figure 1.**
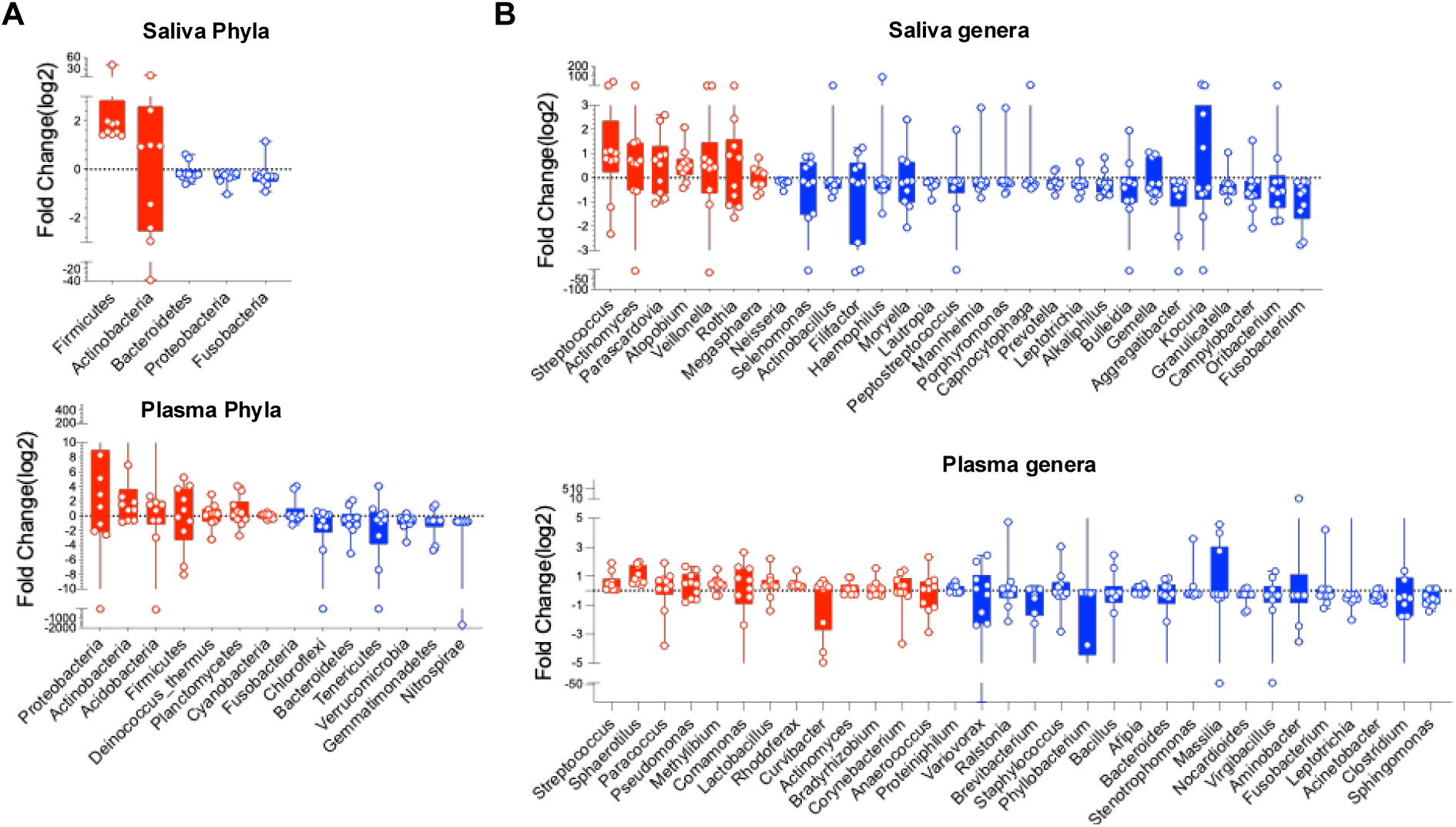
Box plots depict log2 fold changes in bacterial taxa between groups at the (A) phylum and (B) genus levels. Genus column median > 0.007 was selected.

**Supplemental Figure 2.**
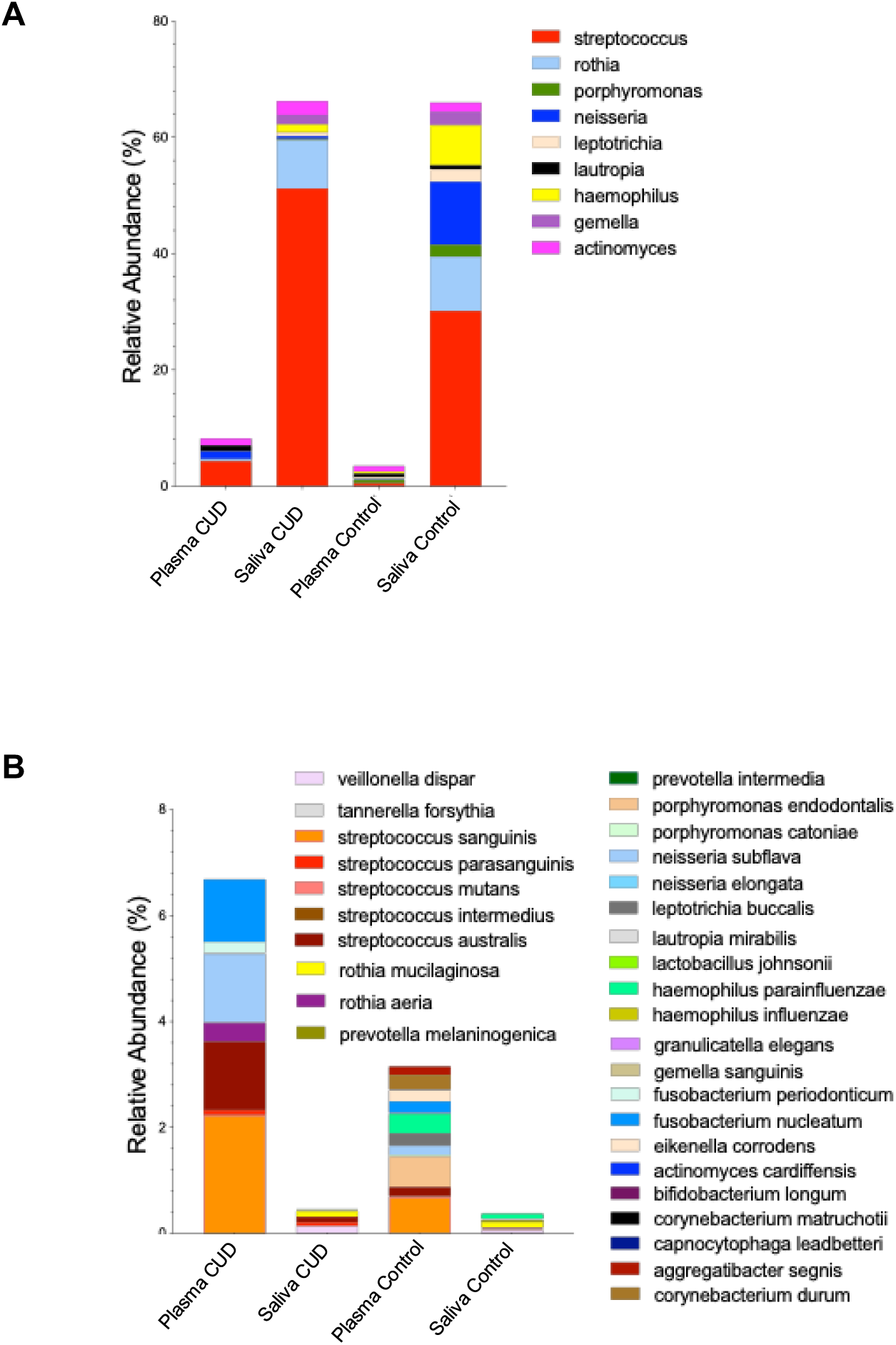
Taxonomic composition of saliva and plasma microbiomes in CUD and controls. Stacked bar plots of bacterial genera (A) and species (B) show the mean enrichment of oral- and plasma-associated taxa.

**Supplemental Figure 3.**
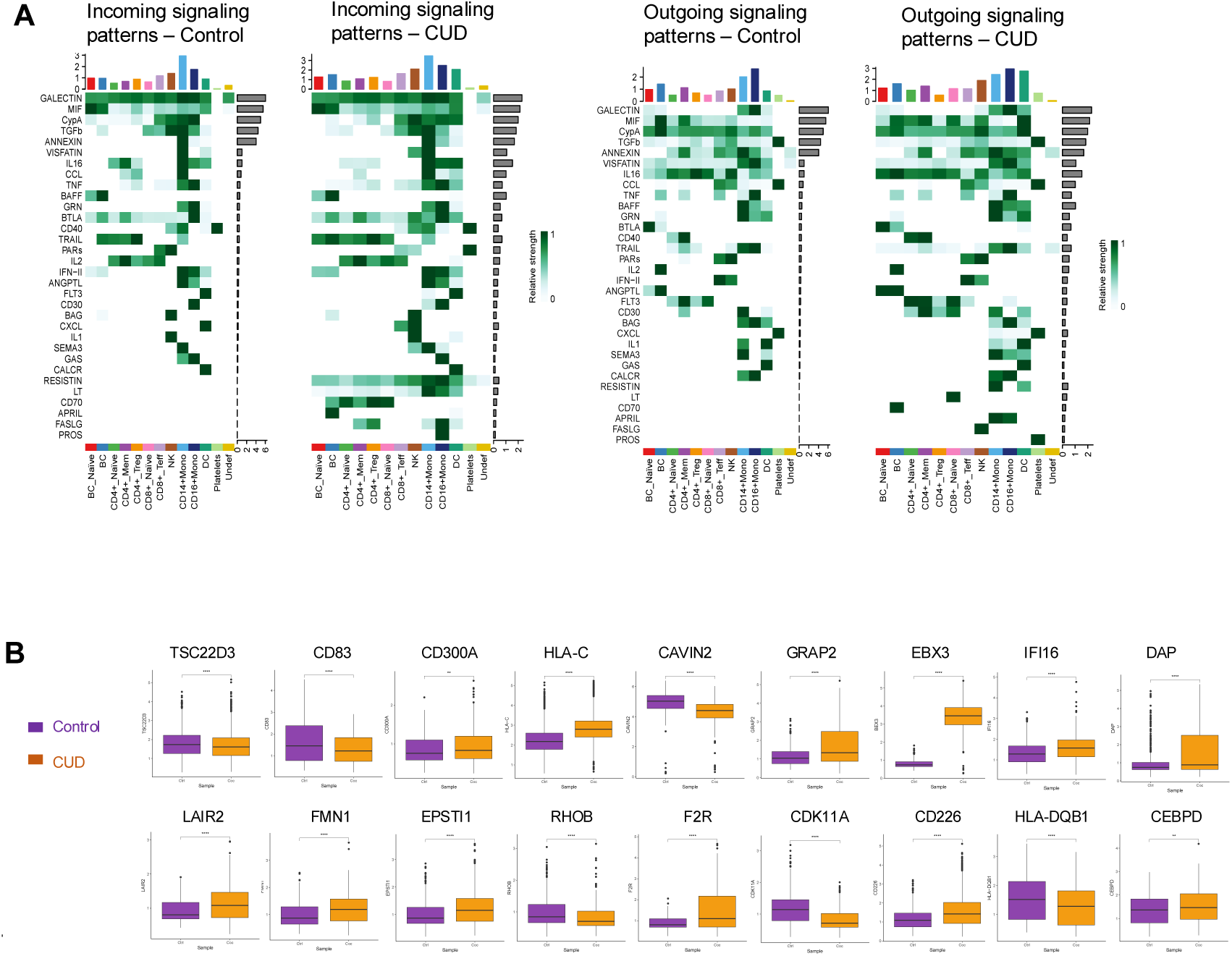
Altered cell–cell communication and gene expression in PBMCs from CUD versus controls. (A) Heatmaps show predicted incoming and outgoing signaling patterns across immune subsets, with CUD samples displaying distinct shifts in ligand–receptor signaling strength compared to controls. (B) Boxplots highlight representative DEGs, including *TSC22D3*, *CD83*, *CD300A*, *HLA-C*, *CAVIN2*, *GRAP2*, *EBX3*, *IFI16*, *DAP*, *LAIR2*, *FMN1*, *EPSTI1*, *RHOB*, *F2R*, *CDK11A*, *CD226*, *HLA-DQB1*, and *CEBPD*, demonstrating significant transcriptional alterations in CUD.

